# A DNA-based optical nanosensor for *in vivo* imaging of acetylcholine in the peripheral nervous system

**DOI:** 10.1101/2020.07.06.189696

**Authors:** Junfei Xia, Hongrong Yang, Michelle Mu, Nicholas Micovic, Kira E. Poskanzer, James R. Monaghan, Heather A. Clark

## Abstract

The ability to monitor the release of neurotransmitters during synaptic transmission would significantly impact the diagnosis and treatment of neurological disease. Here, we present a DNA-based enzymatic nanosensor for quantitative detection of acetylcholine (ACh) in the peripheral nervous system of living mice. ACh nanosensors consist of DNA as a scaffold, acetylcholinesterase as a recognition component, pH-sensitive fluorophores as signal generators, and α-bungarotoxin as a targeting moiety. We demonstrate the utility of the nanosensors in the submandibular ganglia of living mice to sensitively detect ACh ranging from 0.228 μM to 358 μM. In addition, the sensor response upon electrical stimulation of the efferent nerve is dose-dependent, reversible, and we observe a reduction of ~76% in sensor signal upon pharmacological inhibition of ACh release. Equipped with an advanced imaging processing tool, we further spatially resolve ACh signal propagation on the tissue level. Our platform enables sensitive measurement and mapping of ACh transmission in the peripheral nervous system.

## Introduction

The cholinergic system has an important role in nearly every bodily function ranging from muscle contraction, heart rate, gland secretion, inflammation, to cognition and memory. Defects in cholinergic transmission lead to a number of pathologies ranging from Alzheimer’s Disease and Parkinson’s Disease in the central nervous system (CNS)^1, 2^ to Myasthenia Gravis in the peripheral nervous system (PNS)^3^. To understand the many functions of the cholinergic system and the etiology of cholinergic diseases, a longstanding goal has been to develop technologies that allow the sensing and mapping of cholinergic neural communication *in vivo*^4–6^. An obstacle in advancing our understanding of the cholinergic system is the lack of proper techniques for measuring and mapping the spatial and temporal patterns of acetylcholine (ACh) release in organs in real-time with satisfactory sensitivity.

Although there are standard tools to detect ACh in the CNS such as microelectrodes and microdialysis, it is particularly challenging to make measurements in the PNS. ACh acts as a point-to-point neuromodulator in the PNS^7^, which requires precise localization of the probes for spatial resolution and fast response of the probes for temporal information. The size of microelectrodes and microdialysis probes make placement in the PNS difficult, particularly if multiple probes are to be used simultaneously to gain spatial information. Although microelectrodes have unparalleled temporal response, microdialysis can take minutes. Conversely, microdialysis can specifically identify ACh after extraction, but microelectrodes are more suitable for *in vivo* detection of catecholamines than inoxidable ACh^8^. The last decade has witnessed considerable progress in improving microdialysis and microeletrodes for neurotransmitter studies. For example, Zhang *et al.* and Yang *et al.* shortened the sampling time of microdialysis (typically 5-15 min^9^) to only 1-2 min for serotonin analysis via instrumentational upgrades^10–12^, which enabled study of the serotonin network associated with stress, anxiety and circadian rhythms^11, 13^. Song and coworkers achieved high temporal resolution monitoring of ACh with ~5 s intervals in the rat brain by a combination of microdialysis sampling, segmented flow, and direct infusion mass spectrometry^14^. Additionally, many efforts greatly improved the spatial resolution of microelectrodes/fast-scan cyclic voltammetry by minimizing the size of the electrode tip into nanoscale (50 to 400 nm)^15^, even under 20 nm to allow access to single synapses in cultured neurons^16^. New imaging technologies, such as genetically encoded fluorescent sensors (CNiFERs) and G-protein-coupled receptors based sensors with enhanced quantum yield and sensitivity has drawn tremendous attention for *in vivo* imaging of dopamine, norepinephrine and ACh in CNS^6, 17–22^. For the PNS, it would be advantageous to have a sensor that does not require genetic engineering of cells or animals. To meet this goal, we previously reported a nanosensor to detect ACh with a fluorescent reporter, but it lacks the sensitivity to make quantitative measurements in the PNS^23^. Despite these compelling breakthroughs, reports on measuring ACh dynamics in PNS remain scarce, illuminating the difficulty of combining the individual strengths of spatial resolution, temporal resolution, and sensitivity into one measurement tool.

Here, we report a fluorescent ACh-selective nanosensor to image ACh release quantititatively in the PNS and its use in the submandibular ganglion (SMG) of living mice to report real-time, quantitative endogenous ACh release triggered by electrical stimulation of the nerve (Fig. 1a&b). Our design uses double-stranded DNA as the scaffold, acetylcholinesterase (AChE) as the recognition component, and a pH-sensitive fluorophore (pHAb) as the signal reporter (Fig. 1c). Compared to synthetic polymers, using DNA as the sensor scaffold is advantageous because of its ability to form complex structures with a resulting homogeneous size distribution, high yield for ligand attachment to the structure, and exquisite control over positioning of the ligands^24–26^. These properties are important because the sensor mechanism requires that we position the pH-senstive fluorophores in close proximity to AChE to report local pH changes in response to ACh hydrolysis. We also include a second ligand, α-bungarotoxin (BTX), to target the nanosensors to nicotinic ACh receptors^27^. The BTX is additionally labeled with an Alexa 647 fluorophore (BTX-AF647) spectrally-distinct from pHAb and acts as an internal standard, enabling quantitative measurements despite potential variations in sensor number or diffusion in the organ. Additionally, coupled with the Astrocyte Quantitative Analysis (AQuA) program^28^, an advanced imaging processing tool, we further present the propagation of ACh transmission across the whole SMG based on the fluorescence output of the ACh nanosensors.

**Fig. 1.**
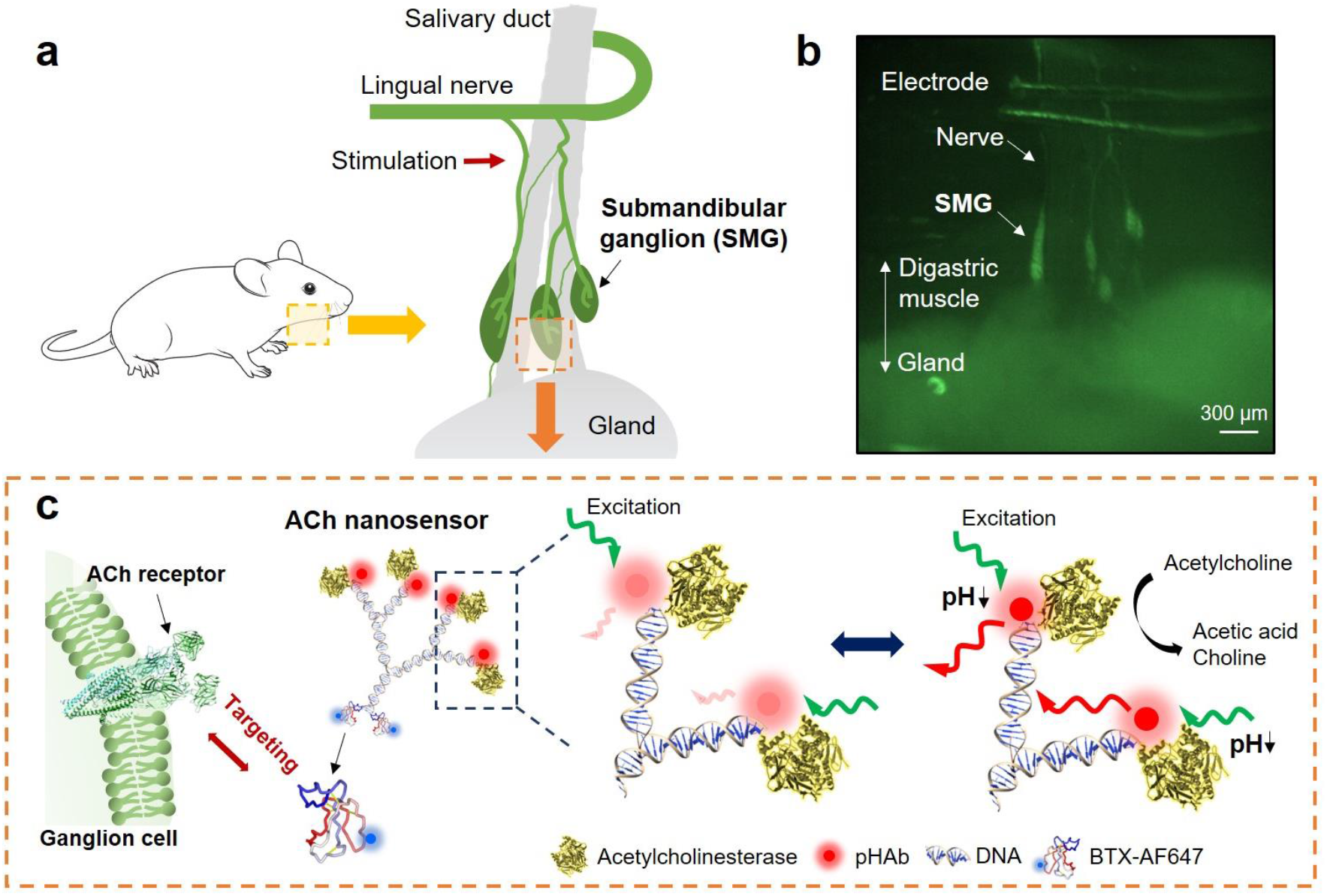
Design of nanosensor for *in vivo* imaging of ACh release. (a) Schematic showing the SMGs located on the salivary duct that connects the digastric muscle to the salivary gland. (b) Image of the SMG in a ChAT^(BAC)^-eGFP transgenic mouse showing GFP expression in presynaptic axons and postsynaptic neurons. Location of the stimulation electrode is indicated. (c) Schematic showing ACh nanosensor structure and mechanism for ACh detection. Nanosensors are targeted to ACh receptors via the conjugated BTX. AChE linked to the DNA scaffold hydrolyzes ACh and lowers local pH through the production of acetic acid. Four pH-sensitive pHAb fluorophores are in close proximity to AChE and increase fluorescent emission in response to ACh hydrolysis. AF647 on the BTX acts as an internal standard to enable quantitative measurements.

## Results

### Fabrication and *in vitro* characterization of ACh nanosensors

The ACh nanosensors consisted of four functional motifs: DNA scaffold, AChE, pHAb fluorophores and BTX-AF647. The dendritic, double-stranded DNA scaffold was self-assembled from five DNA oligonucleotides (*i.e.*, Strands L1 to L5, Supplementary Table 1) and provided 10 anchor points: four for pHAb fluorophores, four for AChE, and two for BTX-AF647 (Fig. 2a). The pHAb fluorophore (3’,6’-bis(4-(4-sulfobutyl)piperazin-1-yl) rhodamine (5’,6’) succinimidyl ester^29^) was selected as the reporter because of its red emission wavelength and “turn on” response when protonated. The pHAb fluorophores were linked to single-stranded DNA (ssDNA) (strands L2, L3, L4) via N-hydroxysuccinimide ester reaction while the other two strands (L1&L5) were functionalized with tetrazine groups for later attachment to BTX (Supplementary Fig. 1). After the five ssDNA oligonucleotides were annealed to form a DNA scaffold, AChE was covalently attached to the DNA scaffold via a maleimide-succinimidyl ester cross linker (maleimide linker) conjugating the primary amine of lysine residues on AChE to the thiol-terminated DNA strands (Fig. 2b). The extremely high catalytic activity of AChE ensures rapid hydrolysis of ACh in a highly specific manner^30^. BTX was chosen as a targeting ligand because it has been widely recognized to bind to nicotinic ACh receptors in a highly selective and efficient manner^31^. A trans-cyclooctene (TCO)-succinimidyl ester cross linker (TCO linker) was conjugated to the surface lysine of BTX-AF647, and further reacted with a tetrazine-terminated DNA via a Click reaction (Fig. 2c).

**Fig. 2.**
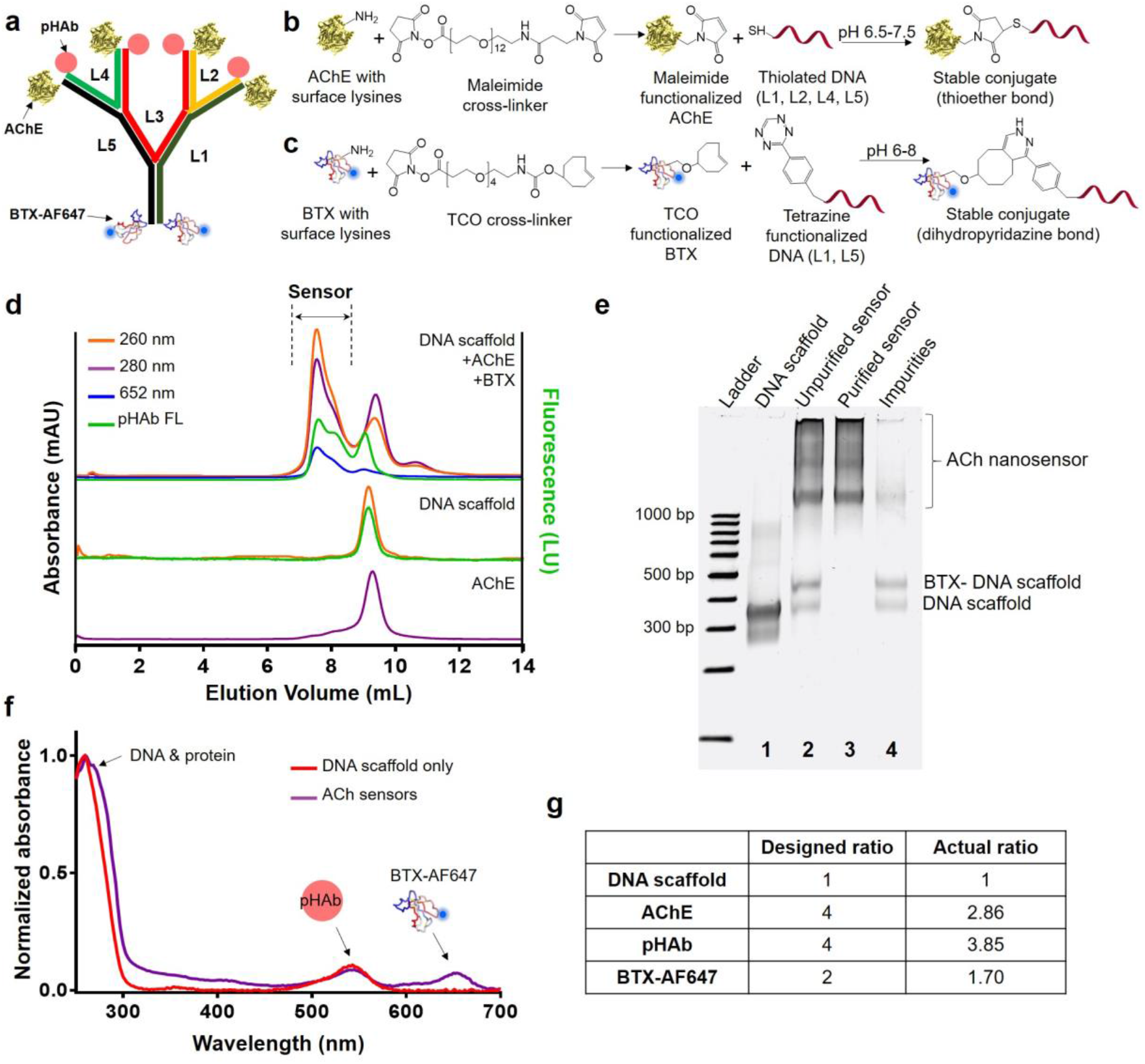
Successful bottom-up assembly and purification of ACh nanosensors. (a) ACh nanosensors consisted of four components: DNA scaffold (five strands, designated L1 to L5), AChE as a recognition motif, pHAb fluorophore as a pH reporter and BTX as a targeting molecule. (b) Conjugation of AChE to DNA scaffold via maleimide-thiol chemistry. (c) Conjugation of BTX to DNA scaffold via TCO-tetrazine click chemistry. (d) Fully-assembled ACh nanosensors were purified by size-exclusion HPLC. Eluted samples from top to bottom: the reaction mixture that contained DNA scaffold, BTX and AChE; DNA scaffold only; AChE only. Three UV-Vis signals and one fluorescent signal were monitored during HPLC elution. Conditions: 150 mM sodium phosphate, 500 mM NaCl (pH=7.2 to 7.4). (e) Native PAGE characterization of the fractions from HPLC in (d) stained with ethidium bromide. Lane 1: DNA scaffold only, Lane 2: Unpurified reaction mixture; Lane 3: Purified ACh nanosensors (main peak); Lane 4: Impurities that contained unreacted AChE, DNA scaffold and BTX-conjugated DNA scaffold (second peak). Unreacted AChE was not shown in gel. (f) UV-Vis spectra of DNA scaffold and ACh nanosensors indicated successful conjugation of two proteins and pHAb fluorophores. (g) Actual conjugation ratios of AChE and BTX on DNA scaffold compared to designed ratios.

Fully-assembled ACh nanosensors were purified by size-exclusion high-performance liquid chromatography (HPLC). A typical chromatogram of the reaction mixture (Fig. 2d) showed well-resolved and reproducible peaks across different batches. ACh nanosensors eluted first (Main peak) due to its high molecular weight, with a retention volume of ~ 7.5 mL, and was identified by three characteristic absorbance spectra (*i.e.*, λ_abs_ = 260 nm for DNA, 280 nm for protein, 652 nm for AF647) and an emission spectrum of the pHAb fluorophore (*i.e.*, λ_ex_=532 nm, λ_em_=560 nm). The following peak with the retention volume of 9.1-9.2 mL included unreacted DNA scaffolds, AChE, and BTX-conjugated DNA scaffolds (Second peak). It was estimated that ACh nanosensors represent approximately 77% of the reaction mixture based on the integration of areas under all detectable peaks from the pHAb channel chromatogram, indicating a high yield in the DNA-protein conjugation. We found that the attachment of BTX to the DNA scaffold alone does not significantly impact the retention time in the Superdex 200 column, evidenced by the approximately identical retention time of BTX-conjugated DNA scaffold in HPLC elution (Data not shown). Note that BTX is a small polypeptide (~8000 Da) compared to the DNA scaffold (~86 kDa) and AChE (280 kDa). Fractions from the main and second peak were collected and further characterized by native polyacrylamide gel electrophoresis (PAGE). Compared to the DNA scaffold, several bands with lower mobility were identified likely due to variable numbers of AChE (from 1 to 4) conjugated to the DNA scaffold (Fig. 2e). Bands that presumptively contain ACh nanosensors displayed overlapping fluorescence of pHAb and AF647, demonstrating structural integrity of ACh nanosensors (Supplementary Fig. 2). The purified ACh nanosensors exhibited peak absorbances around 260-280 nm arising from the DNA and protein, 532 nm from pHAb fluorophores, and 647 nm from AF647 (Fig. 2f). To calculate the ratio of conjugated AChE and BTX to DNA scaffold, we estimated concentration of BTX-AF647 ([BTX]) by measuring the absorbance at its maximum absorbance of 652 nm. We then estimated extinction coefficients (*i.e.*, ∊) of the DNA scaffold, AChE, and BTX at 260 nm and 280 nm by measuring the absorbance of pure DNA or protein samples.

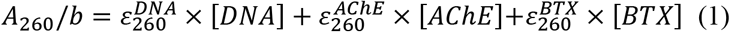

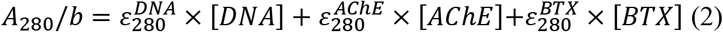

Based on the above Beer-Lambert Law calculations, the conjugation ratios of AChE, BTX, and pHAb to DNA scaffold were determined to be 2.86, 1.70 and 3.85 (Fig. 2g). The conjugation efficiency of BTX and pHAb reached 85% and 96%, respectively. The slightly lower conjugation efficiency of AChE (*i.e.*, 71%) may be attributed to steric hindrance during the DNA-enzyme reaction. The agreement between measured and designed ratios supports the high predictability of the DNA nanostructure self-assembly. Compared to our previous version with an estimated size of ~130 nm^23^, the current ACh nanosensor had an overall size of 10-20 nm confirmed by both structural simulations and atomic force microscopy (Supplementary Fig. 3). It is worth noting that due to the inherent flexibility of DNA structures^32, 33^, the hydrodynamic size of ACh nanosensor is even smaller, making it suitable for probing small biological systems.

### ACh nanosensors in SMG

The SMG parasympathetic ganglia that innervate the salivary glands was chosen to validate our nanosensor because 1) it is relatively accessible for optical imaging, 2) it is a well-established model of ACh release, and 3) the SMG contains large neurons with no dentrites that are mainly innervated by a single cholinergic fiber that activate nitotinic ACh receptors on the cell soma^34, 35^. To expose the SMG, we adapted a protocol for time-lapse imaging of interneuronal synapses in SMG from previous literature^36, 37^. We first made a small incision on the neck and then placed an insulated metal platform under the salivary ducts to lift and stabilize the SMG to be imaged with a water-immersion objective. The flat metallic support greatly reduced respiration-induced movements which would cause defocusing and signal fluctuations during imaging. Several SMGs can be identified along the salivary ducts by removing the nearby connective tissues (Supplementary Fig. 4 a&b). *Ex vivo* staining of freshly isolated SMG with BTX-AF647 showed the expected punctate staining pattern across the ganglion, confirming expression of nicotinic ACh receptors in SMG neurons (Supplementary Fig. 5).

ACh nanosensors were administered *in vivo* to the SMG by local microinjection. Fig. 3a shows the setup for microinjection: a pulled glass needle filled with nanosensors was used for injection, by rapid filling of extracellular space in the SMG, which can be observed as fluorescent rings surrounding each ganglion cell after injection (Supplementary Video 1). Injection was performed using a a widefield microscope with low-magnification (4× objective) and dual imaging of pHAb and AF647 channels (Fig. 3a) (detailed in Methods). Some ACh nanosensors were found along the nerve tract that entered and exited the SMG (Supplementary Fig. 4c). After extensive buffer rinsing, a significant amount of ACh nanosensors remained inside the SMG (Fig. 3b). Overlap fluorescence of BTX-AF647 and pHAb indicated that the ACh nanosensors maintained structural integrity in the tissue. In contrast, DNA structures that did not have the targeting ligands (*i.e.*, BTX) were barely visible inside the tissue (Fig. 3c). Confocal microscopy further confirmed that the punctate staining pattern of AF647 and pHAb were primarily localized on the neuronal cell soma (Fig. 3d). The consistent staining patterns of ACh nanosensors in the SMG *in vivo* along with the *ex vivo* staining of the SMG with BTX-AF647 (Supplementary Fig. 5) support that ACh nanosensors bind to nicotinic ACh receptors via conjugated BTX.

**Fig. 3.**
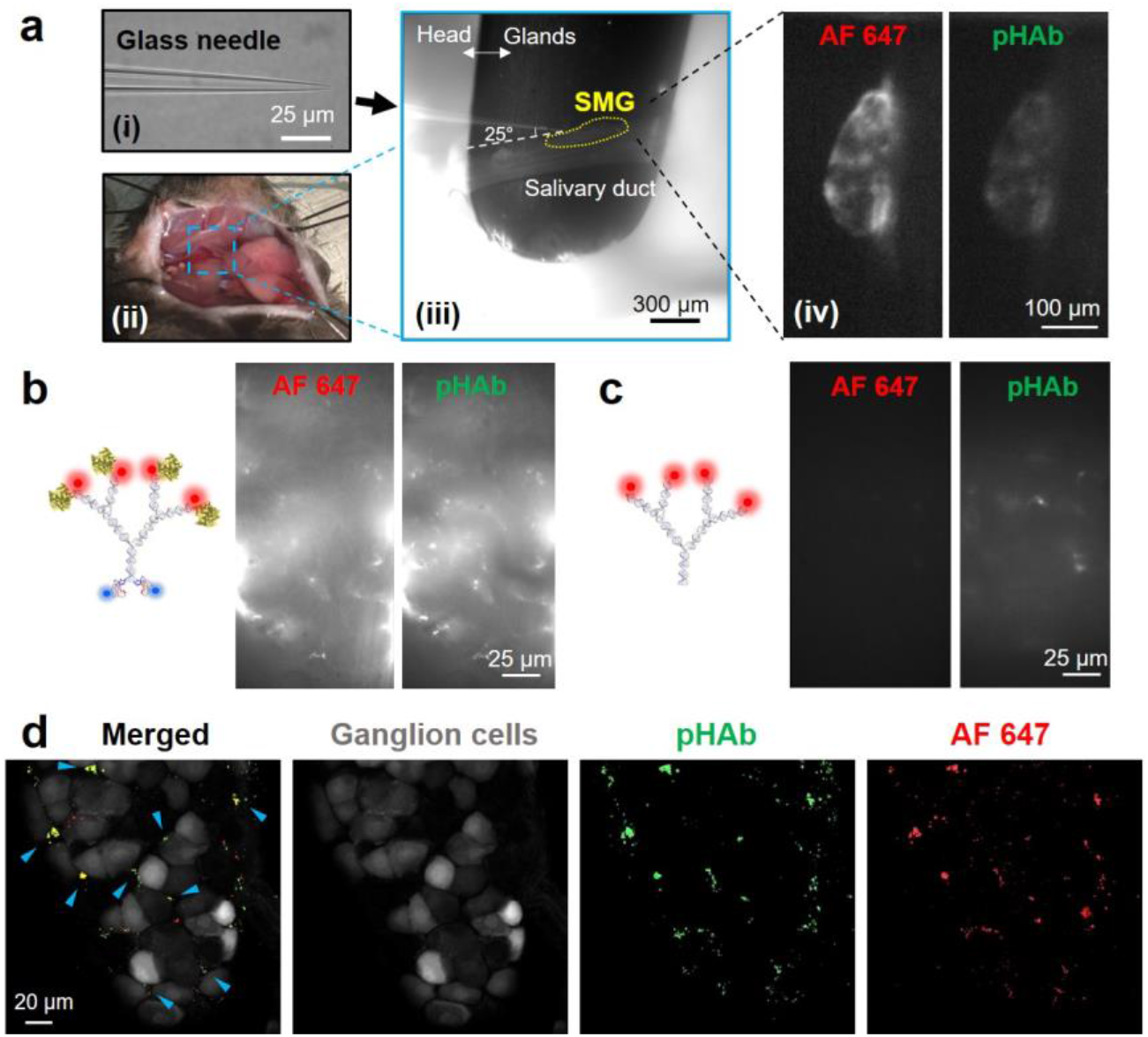
ACh nanosensors in SMG. (a) Microinjection of the ACh nanosensors into an SMG of a living mouse via (i) a glass needle. Scale bar: 25 μm. (ii)&(iii) The SMG on a salivary duct was exposed by surgical incision and moving the salivary gland away from the duct. (iii) Scale bar: 300 μm. (iv) ACh nanosensor-stained SMG observed under 4× objective. Scale bar: 100 μm. (b) ACh nanosensors and (c) DNA structures that do not have BTX-AF647 were washed away after injection. Scale bar: 25 μm. (d) Confocal imaging of the nanosensor-stained SMG in a ChAT^(BAC)^-eGFP transgenic mouse after removal and fixation. Grey: ganglion cells; Green: pHAb fluorophores; Red: AF647 fluorophores. Yellow (highlighted by blue arrows) indicates the overlap of the pHAb and AF647 fluorophores. Scale bar: 20 μm.

### *Ex vivo* calibration of the ACh nanosensors

We next measured the response of the ACh nanosensors *ex vivo* using exogenous ACh to build a calibration curve. The SMG was microinjected *in vivo* with nanosensors, isolated, and then imaged by positioning the SMG under a metal harp in a bath imaging chamber (Fig. 4a) as we changed the ACh concentration in the surrounding bath solution. Qualitatively, we observed a significant rise in fluorescence intensity as the ACh concentration increased stepwise from 0.71 nM to 7.14 mM (Supplementary Video 2), as predicted. The initial rates of change in pHAb fluorescence intensity were fitted and analyzed by the Michaelis-Menten equation (Fig. 4b). From the curve, we see that the the half-maximum response (EC50) to acetylcholine was 49.41 ± 7.66 μM and upper limit of detection was 358.01 μM. To determine the lower limit of detection (LLOD), experimental data in the lower concentration range (*i.e.*, 0.71 nM to 14.28 μM) was grouped into two linear regions with the two regression lines crossing at a value of [ACh]=228 nM. The theoretical nanosensor response at LLOD of 0.72% in unit recording time (*i.e.*, increase rate 1.3%/s is derived from Michaelis-Menten equation and the recording internal is 0.55 s) was significantly higher than the background noise (*i.e.*, 0.088%±0.039%). The signal-to-noise ratio of the LLOD was 8.18, which provided an estimate of [ACh]=228 nM as the LLOD. It has been estimated that the concentration of ACh ranges from low-to mid-micromolar *in vivo*, which lies within the detection range of our ACh nanosensor^38–40^. Concerns may arise regarding whether the ACh nanosensor’s response can be influenced by the amount of nanosensors present in the SMG. Thus, we separately normalized the initial rate (*i.e.*, V_sensor_, ΔF/F per second, %/s) with the unit initial rate (*i.e.*, V_unit sensor_, ΔF/F per second per nanosensor, %/(s·nM)). Specifically, the nanosensor concentration in the region of interest (ROI) was calculated based on the fluorescence intensity of AF647 (internal standard) via a linear calibration curve derived from ACh nanosensors of known concentrations. The normalized unit initial rate V_unit sensor_ was obtained by dividing the original V_sensor_ in Fig. 4c by the as-calculated nanosensor concentration (nM). The V_unit sensor_-[ACh] relationship was further analyzed by the Michaelis-Menten equation (Supplementary Fig. 6), which showed an almost identical curve to Fig. 4c with an EC50 value of 50.35 ± 6.76 μM. A double reciprocal plot was obtained from the fitted Michaelis-Menten equation as Equation (3) and used to quantify ACh concentrations in the *in vivo* experiments.

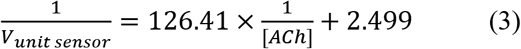

**Fig. 4.**
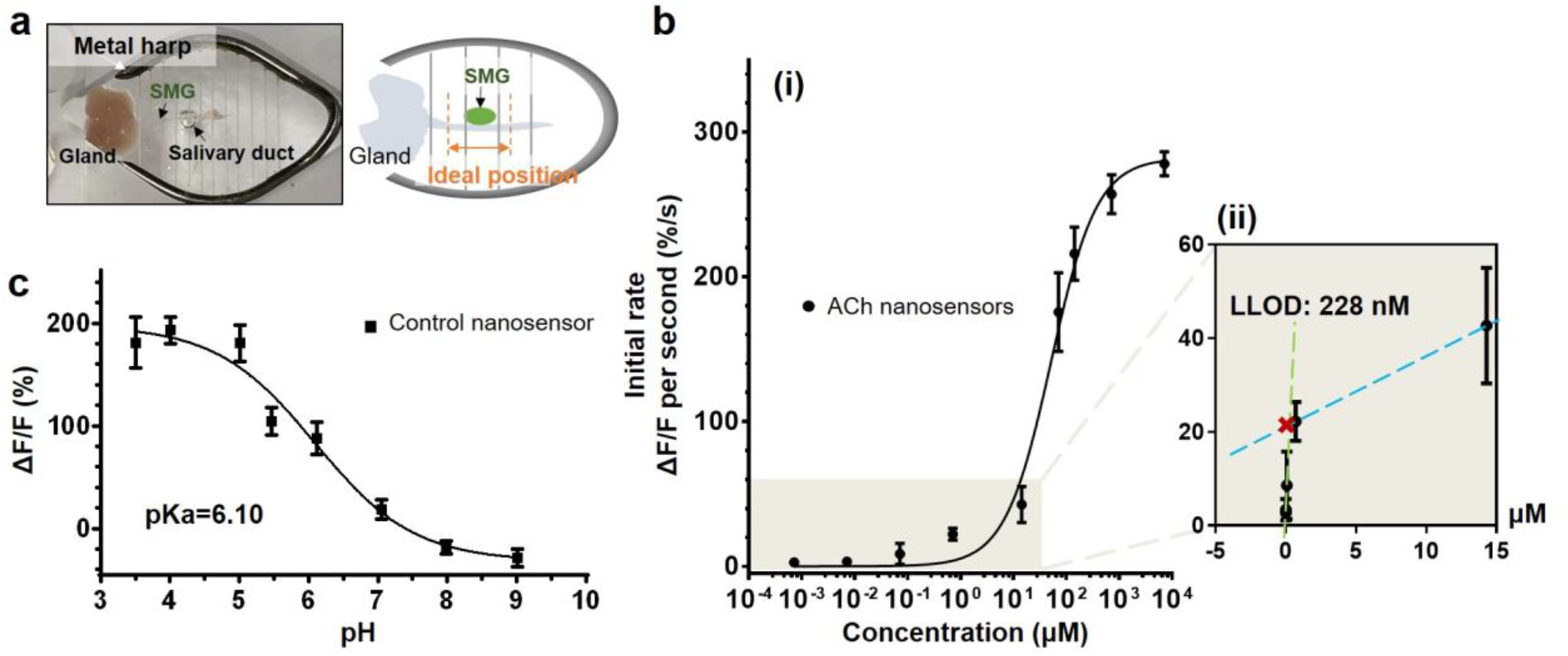
*Ex vivo* calibration of ACh nanosensors. (a) Set-up for the *ex vivo* calibration of the control and ACh nanosensors. The SMG was isolated immediately following the microinjection of the nanosensors into the tissue, and later positioned under a metal harp in a bath imaging chamber for imaging. (b) The initial rates of the sensor response to exogenous ACh solutions of different concentrations. The sigmoidal nonlinear curve was fit by the Michaelis-Menten equation with the constants *K*_*m*_=49.41 μM (EC50). (ii) expanded view of the region denoted in grey in (i). The data in the lower concentration range (up to 14.28 μM) were grouped into two linear regions (i.e., green and orange lines); the two dashed lines intersect at [ACh]=228 nM, corresponding to the LLOD. Error bars denote S.D. resulting from triplicate experiments. (c) Fluorescence change of control nanosensors under different pH levels ranging from 3.5 to 9.

As with many optical probes for cellular studies^41–43^, our nanosensors are dependent on pH and could potentially respond to endogenous changes in pH not associated with the sensing mechanism. As a control for measuring pH, we fabricated a nanosensor with only pHAb fluorophores and BTX on the DNA scaffold (no enzyme, termed “control nanosensor”). We calibrated the response of the control nanosensor towards varying pH levels in an *ex vivo* tissue preparation. The signal change from pH 7 to 5 ranged from 18.50 ± 9.74 % to 180.43 ± 17.67 % with a pK_a_ of 6.10 (Fig. 4c), in accordance with the pK_a_ value of 6.2 representing free pHAb fluorophores (Supplementary Fig. 7a). The large signal change in mildly acidic conditions is a property that favors detection of ACh with our proposed sensing mechanism that generates a localized acidic environment in the vicinity of the enzymes.

### *In vivo* imaging of ACh release

ACh nanosensors were microinjected into the SMG as previously shown in Fig. 3a. Electrical stimulation of the lingual nerve was performed (3V, 10 Hz, 10 ms duration, 100 pulses) to evoke release of ACh into the SMG (Fig. 5a; Supplementary Fig. 4a&b)^44^. Upon the stimulation of the lingual nerve, fluorescence intensity rapidly increased 3.7 % by 15 seconds and slowly recovered to baseline within 2 mins (Fig. 5b). Stimulation evoked a steady increase in signal over ~15s, likely indicating persistent pre-synaptic firing. In contrast, we observed a minimal reponse when control nanosensors were tested, demonstrating that the influence of intrinsic pH fluctuation is negligible. However, upon stimulation with a higher 10V voltage, control nanosensors responded to ~50% of the ACh nanosensor signal intensity (Supplementary Fig. 8), suggesting higher stimulation induces a pH change, possibly due to cellular damage or endogenous enzyme. Varying the electrode stimulation parameters led to a dose-dependent ACh nanosensor response (V_sensor_) with 3V stimulation increasing 0.1886%/s, 5V increasing 3.24%/s, and 10V increasing 11.35%/s, all collected at the same ROI in the ganglia (Fig. 5c). Similarly, increasing the number of pulses also led to a higher ACh nanosensor response (Fig. 5d). Reversibility of the sensor was demonstrated by three repeated stimulations (10V, 10 Hz, 10 ms duration, 10 pulses), showing repeatable rates of 0.436%/s, 0.417%/s, and 0.431%/s in sequence (Fig. 5e), which was confirmed with *in vitro* experiments (Supplemental Figure 6b).

**Fig. 5.**
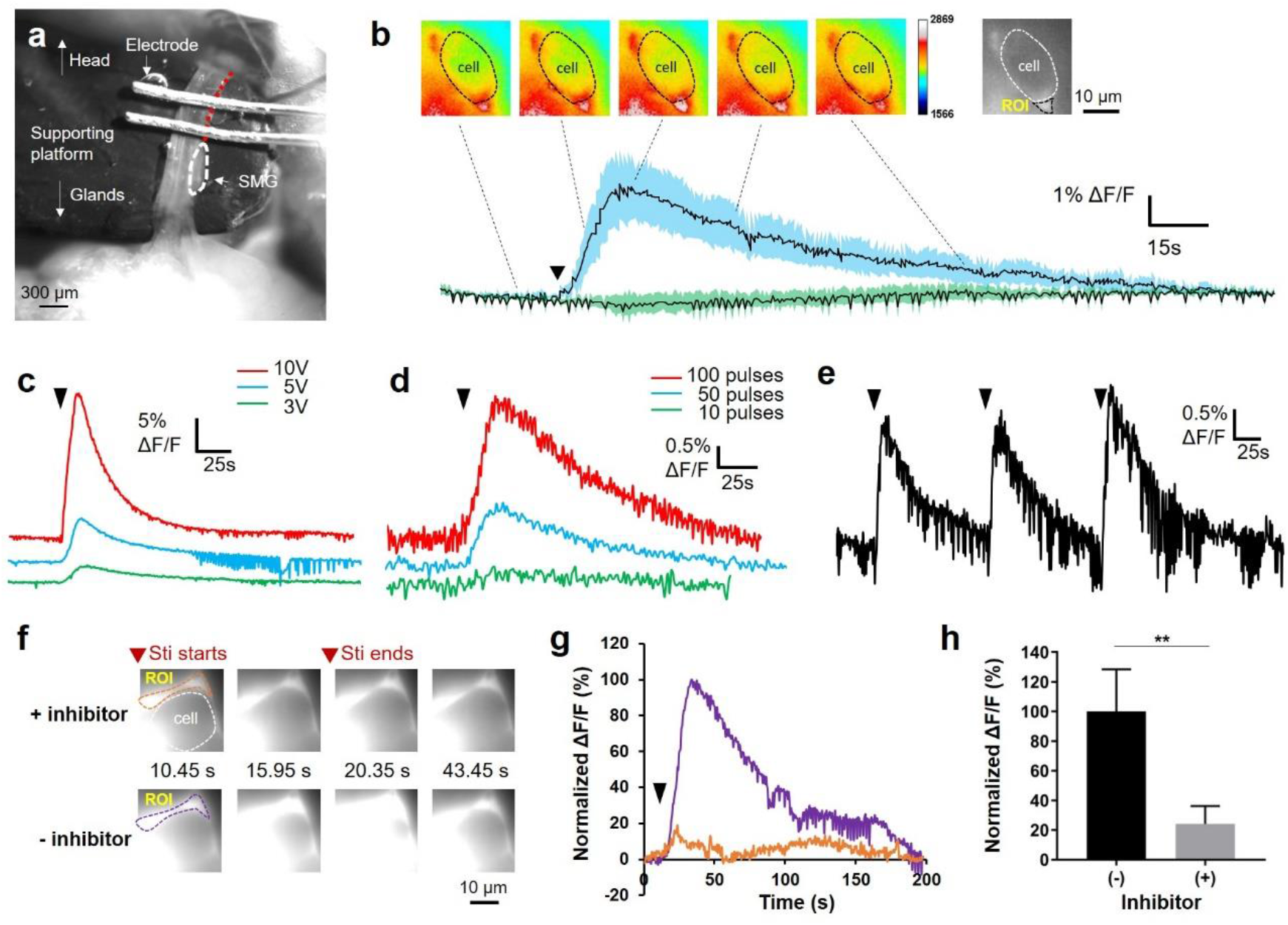
*in vivo* imaging of ACh release. (a) Bright field image showing the setup of electrical stimulation. Red dashed line indicates segments of nerves being stimulated. Scale bar: 300 μm. (b) ΔF/F of fluorescence intensity from the pHAb channel in response to electrical stimulation (indicated by black arrow). The panel of time-lapse fluorescence images display a typical ACh nanosensor-stained, single ganglion cell used for plotting. The green curve represents data collected from ganglion cells stained with the control nanosensors. Data was averaged from 3 independent experiments. Scale bar: 10 μm. (c) Plot of ΔF/F in the same ROI in response to the electrical stimulation with different voltage amplitudes (i.e., 3V, 5V, and 10V). Other simulation parameters: 10 Hz, 10 ms duration, 100 pulses. (d) Plot of ΔF/F in the same ROI in response to the electrical stimulation with different numbers of pulses (i.e., 100, 50, 10). Other stimulation parameters: 10 Hz, 10 ms duration, 3V. (e) Plot of ΔF/F in the same ROI of 3 repeated electrical stimulations. Stimulation parameters: 10V, 10 Hz, 10 ms duration, 10 pulses. (f)-(h) ACh nanosensor’s response to the ACh release with or without vesamicol, an ACh inhibitor. (f) Time-lapse fluorescence images showing the real-time fluorescence readout before and after washing away the vesamicol. Scale bar: 10 μm. (g) ΔF/F of fluorescence intensity from pHAb channel plotted from ROI (purple and orange dashed line) in (f). (h) The peak value of ACh nanosensor response from four independent inhibitor experiments. By the paired t-test, a significant difference (^**^p < 0.01) was found between the vesamicol treated and untreated groups for all four trials. Fluorescence intensity was normalized at the maximum value in (g) and (h). Stimulation parameters for drug experiments: 3V, 10Hz, 10ms duration, 100 pulses. Black triangles indicate the start of the stimulation.

Next, we quantified ACh concentration using the established calibration curve. In Fig. 5c, The V_unit sensor_ values for electrical stimulation of 3V, 5V, 10V were 0.0214%/(nM·s), 0.0711%/(nM·s), and 0.2448%/(nM·s), respectively. Using Equation (3), we estimated the endogenous ACh concentrations to be 2.75 μM at 3V, 10.49 μM at 5V, and 76.51 μM at 10V. Note that calculated values are above the LLOD with a high signal-to-noise ratio.

As a second confirmation of the response to ACh, an anti-cholinergic drug, vesamicol, was introduced to temporarily block the release of ACh. Vesamicol inhibits ACh release by inhibiting ACh uptake into synaptic vesicles in pre-synaptic neurons^45^. We incubated the nanosensor-stained SMG with vesamicol for 15 min and recorded the sensor’s response with electrical stimulation at 3V. We then washed away the vesamicol and stimulated the lingual nerve under the same stimulation conditions. The ACh nanosensors reported only 17% response with the drug compared to after the drug was washed from the sample (Fig 5f and g). Four independent inhibitor treatments affirmed that vesamicol can considerably suppress ACh nanosensors’ readout by an average of 76.59 ± 11.34 % (Fig. 5h), serving as a strong evidence that the ACh nanosensor signal indeed stems from endogenous ACh release.

### ACh nanosensor response reflects physiological ACh propagation

We observed that lingual nerve stimulation induced a complex spatiotemporal response across the SMG. To quantify this complex response, we applied the AQuA software which can accurately quantify irregular fluorescence dynamics^28^. In AQuA, coordinated fluorescence dynamics are classified as “events”, and these events vary in area, duration, amplitude, and propagation. As such, no predetermined ROIs are selected, for a less biased method of image analysis. AQuA is particularly powerful in the analysis of fluorescence dynamics that are irregular, heterogeneous, and/or propagative. Here, we used AQuA to analyze the fluctuations in intensity of the individual pixels within SMG. To compare the temporal ACh nanosensor response calculation between AQuA to our original ROI-based analysis, we compared the fluorescence dynamics derived from AQuA software to the ROI-based analysis at the same regions that were automatically defined by AQuA software, and found that they reported similar dynamics (Supplementary Fig. 9). We next applied AQuA to quantify the spatial propagation of the fluorescent signal across the ganglion. Since the SMG is a well-established model of ACh release^34, 35^ and we stimulated preganglionic axons based on established methods^36, 37^, we expected to observe rostral-to-caudal ACh transmission. The spatiotemporal AQuA analysis of nanosensor response following 8 V stimulation across the SMG validated our prediction (Fig. 6a and Supplementary Video 3). Stimulation of 8V generated an increase in fluorescence intensity at the rostral-most region peaking at t=22.00 s, 1.65 s after the last stimulation. This signal propagated caudally by t=26.95 s, and further traveled laterally to the light purple region where decay begins at t=44.55 s. All detected events start to decay after t=44.55 s, 24.2 s after the last stimulation (Fig. 6b). All evoked nanosensor responses in Fig. 6a climbed to the peak around ~10-15 s, an observation consistent with Fig. 5b-e. To support the finding that ACh nanosensor responses reflect physiological ACh propagation, we also conducted the same analysis on the same experiment with a lower stimulation voltage of 3 V, and observed similar rostral-to-caudal propagation directionality, but with lower amplitude (Supplementary Video 4), consistent with the dose-dependent sensor response in Fig. 5c. It is worth noting that previous work focused more on single cell recording^46^, ion-channel^47^ and receptor recycling^48^ of ganglion cells; we instead demonstrated the spatial direction of ACh transmission in the SMG for the first time.

**Fig. 6.**
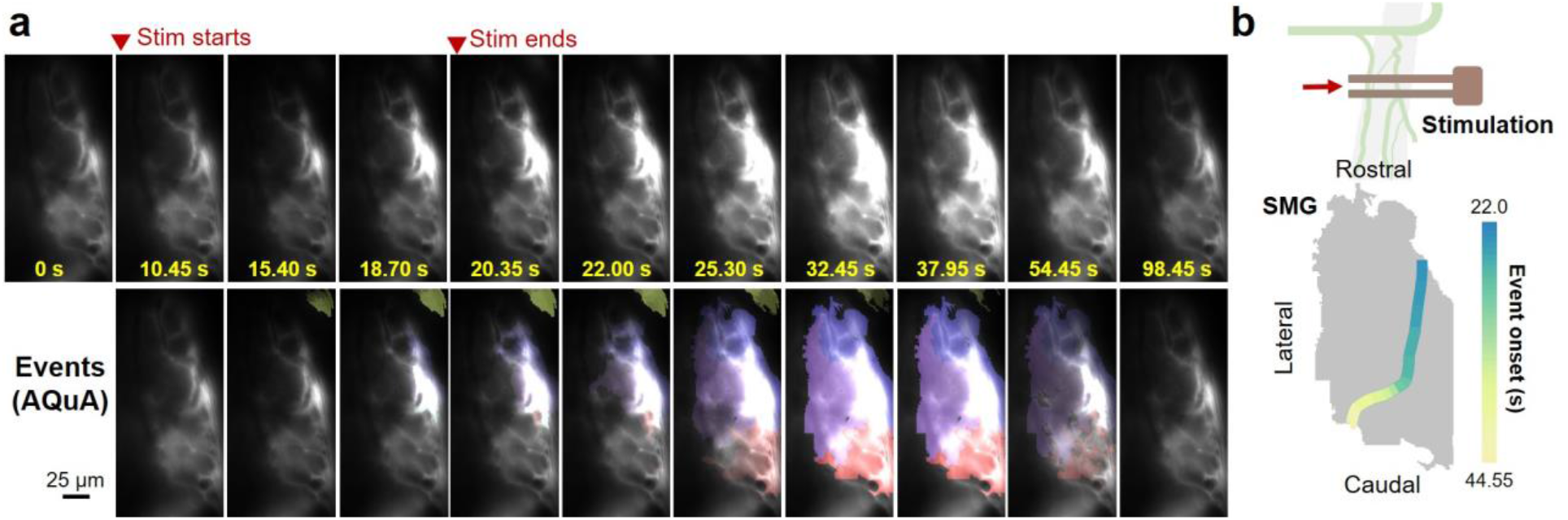
AQuA program resolves the ACh propagation directionality. (a) Time-lapse images (upper panel) exhibit a whole SMG injected with the ACh nanosensors under electrical stimulation. Bottom panel shows the same series of images with the AQuA-detected events. Scale bar: 25 μm. (b) *in vivo* ACh transmission propagates with specific directionality in (a). Stimulation parameters: 8V, 10 Hz, 10 ms duration, 100 pulses.

## Discussion

In this study, we used a bottom-up modular design and assembly process to develop a DNA-based enzymatic nanosensor for detecting ACh transmission in the PNS. The DNA scaffold plays a key role in the accurately positioning the functional motifs AChE and pHAb in close proximity for sensitive detection of the pH drop caused by ACh hydrolysis. It should be mentioned that the ACh nanosensor is highly versatile and can be easily re-purposed for other applications or imaging in different regions of the brian and body. In addition, the AChE and pH-sensitive fluorophores can be substituted with other functional motifs to target other neurotransmitters (e.g., tyrosinase and oxygen-sensitive fluorophores for sensing dopamine)^49^. The targeting ligand, BTX, can be replaced by other biomolecules for targeting other receptors of interest. Additionally, the structure of the DNA scaffold can be easily adjusted to accommodate customized needs and prevent degradation *in vivo*. In this work, the size of the ACh nanosensor is specifically designed to be 10-20 nm to facilitate localization within complex, heterogeneous tissues. We chose nicotinic ACh receptors as the binding target because it is used broadly across the body including the CNS and sympathetic, parasympathetic, and somatic PNS and because of its well-studied binding affinity to BTX^27, 31^. Successful employment of the same receptors as the targeting region is also demonstrated in CNiFER sensors^6, 50^. Concerns may arise regarding whether the binding between BTX and ACh receptors would block normal ACh signal transmission and negatively affect or harm the imaging subject. We hypothesize that a low percentage of ACh receptors would be occupied by the ACh nanosensors so a substantial number of ACh receptors in the synapses would still remain available. The availability of additional ACh receptors is also supported by the regeneration of membrane-bound ACh receptors in SMG via continuously removing/inserting receptors from/to the plasma membrane ^51^.

A major accomplishment in this work is the successful imaging of endogenous ACh release in the SMG, as a proof-of-concept tissue in the PNS, and the tracing of ACh transmission on the tissue level. Two principal factors support that the detected signal originates from endogenously released ACh: (1) electrical pulses trigger dose-dependent and reversible responses from the ACh nanosensor, and (2) the signal can be significantly recovered after removal of vesamicol, a pre-synaptic ACh inhibitor. The quantification of endogenously released ACh is achieved via establishing an *ex vivo* calibration curve. As reported, ACh concentrations lie in the range of low to mid-micromolar predominantly in the CNS, whereas few estimates have been made in the PNS except for the neuromuscular junctions. In contrast to single-point detection tools, such as electrodes which are ineffective against spatial mapping, the benefit to an imaging probe is the ability to attain spatial information in addition to temporal especially when the signal is irregular and propagative. Our data provides new insights into measuring ACh concentration in the PNS and spatial mapping of ACh signal transmission on the tissue level, a significant advance in resolving cholinergic transmission in the PNS. The propagation directionality derived from advanced algorithmic image processing facilitates spatiotemporal mapping of cholinergic transmission. The algorithm can be expanded to 3D imaging of neurotransmitter dynamics using confocal and light sheet microscopy^52^. With the evolution in machine-learning-based image processing and advancement in high-resolution volumetric imaging, we envision this sensing platform will serve as a powerful tool to elucidate the underlying mechanisms in cholinergic circuitry.

As with any tool, there are issues to be addressed in future designs. First, the DNA scaffold was designed for the environment of the SMG and could degrade in the presence of degradative enzymes in the bloodstream. The next stage of development will focus on the scaffold by using enzyme-resistant nucleotides or shielding the sensor with polyethylene glycol (PEG). The fluorophores could be substituted to minimize photobleaching or use near-infrared or two-photon reporters for ideal tissue imaging. In summary, we show a biochemical nanosensor that rapidly responds to endogenous ACh release in a dose-dependent and reversible manner, and exhibits significantly improved spatial flexibility that enables mapping of ACh signal transmission on the whole tissue level. This work sheds light on a new imaging modality that consolidates sensitivity, temporal and spatial resolution, and comprehensively resolves ACh transmission in the PNS.

## Methods

### Design and fabrication of the ACh nanosensor

The structure of the DNA scaffold was designed to accommodate four AChE enzymes (*electrophorus electricus,* Millipore-Sigma), four pHAb fluorophores (Amine-reactive, Promega) and two BTX molecules (AF 647 conjugated, Thermo fisher) while minimizing the overall size. Preliminary sequences were obtained by a random sequence generator^53^ to anneal into a dendritic secondary structure with four branches and a focal point as confirmed by RNAfold minimum free energy calculations^54^ and Nanoengineer-1 Program (Molecular Dynamics Studio). Single-stranded DNA oligonucleotides with amine or thiol functional groups were purchased from Sigma. Before annealing the DNA scaffold, pHAb dyes were conjugated to DNA strands (*i.e.*, L2, L3, L4) via NHS-ester reactions (Supplementary Fig. 2). The same reaction was also employed to modify two amine groups in strands L1 and L5 with tetrazine groups via tetrazine linkers (Tetrazine-PEG5-NHS ester, Broadpharm) (Supplementary Fig. 2), which were later used to conjugate BTX. Reactions were conducted in 0.1 M NaHCO_3_ solution (pH=~8.1-8.2) at a DNA concentration of 100-150 μM. Modified single strands (L1 to L5) were firstly separated from excess pHAb dyes and tetrazine cross linkers by 2 rounds of ethanol extraction, and further purified by illustra NAP-10 columns (GE Health) via gel filtration, followed by overnight lyophilization (Labconco).

The DNA scaffold was self-assembled by incubating 5 DNA oligonucleotides following a temperature gradient from 95 °C to 4 °C for 6 h in a thermocycler (Mastercycler^®^ nexus X2, Eppendorf) in Tris-EDTA buffer (TE buffer, 10mM tris and 1mM EDTA). Misfolded DNA scaffolds were removed by a size-exclusion column (Superdex 200 increase 10/300 GL column, GE Health) via HPLC (Infinity 1260, Agilent) using phosphate buffer (500 mM NaCl) as the eluent (Supplementary Fig. 10a). The correct fraction was collected and resuspended in TE buffer via buffer exchange. The fully assembled DNA scaffold can be stored in a −20 °C freezer for up to 6 months. Two proteins were attached to the DNA scaffold after the annealing process. Before enzyme modification, we firstly employed size-exclusion HPLC to purify AChE since the commercially-available AChE (from eel extract) is not of 100% purity (Supplementary Fig. 10b). The correct fraction was collected for further modification with maleimide functional groups. For protein functionalization, 30 to 50 μM of AChE was swirled with maleimide linkers (maleimide-PEG_12_-NHS ester, Pierce SM(PEG)_12_ Thermo Scientific) for 2 h before being purified by 100 k Amicon centrifugal filter 5 times. BTX was maintained at ~500 μM when reacting with TCO linker (TCO-PEG_4_-NHS ester, Click Chemistry Tools). In the aforementioned protein-linker reactions, either AChE and BTX solution was incubated with corresponding cross linkers at a molar ratio of 1:20 to keep the balance between linkage efficiency and protein activity. Assembled DNA scaffolds were pretreated with TCEP for 1.5 h at RT to cleave the disulfide bonds and then removed from excess TCEP via Amicon centrifugal filters (30k cut-off) prior to enzyme conjugation. Maleimide-functionalized AChE was later vortexed with the DNA scaffold at a molar ratio of 5:1 for 2 h. Afterwards, TCO-functionalized BTX was added to the reaction mixture at a BTX to DNA molar ratio of 20:1, allowing a TCO-tetrazine ligation reaction between BTX and the DNA scaffold for another 2 h. The unreacted thiol groups were later capped by ethylmaleimide (Fisher Chemical). The final reaction mixture was vortexed for 30 min before transferring to a 4 °C fridge for overnight incubation.

After protein-DNA conjugation, ACh nanosensors that contain DNA scaffold, AChE and BTX were separated from other impurities by a size-exclusion column (Superdex 200 increase 10/300 GL column, GE Health) via HPLC (Infinity 1260, Agilent). The elution solution was 0.15 M phosphate buffer with NaCl concentration of 500 mM. Three UV signals (i.e., 260 nm and 280 nm for DNA and protein absorbance, 652 nm for AF 647) and one fluorescence signal (i.e., Ex=532 nm, Em=560 nm for pHAb) were monitored during elution. After purification, ACh nanosensors were concentrated by 100k Amicon^®^ ultra centrifugal filters, and then aliquoted for long-term storage at −20 °C.

### Enzyme assay

The enzymatic activity of AChE was calculated based on the Ellman Esterase Assay^55^. The substrate solution was prepared by mixing 100 μL of Ellman’s reagent (DTNB, 5,5’-dithio-bis-[2-nitrobenzoic acid], 3.96 mg/mL), 40 μL of 75 mM acetylthiocholine iodide (Millipore-Sigma), and 3 mL of 100 mM sodium phosphate buffer (pH=7.0-8.0). 200 μL of the substrate solution was firstly added to a 96-well plate. After 40 μL of AChE/ACh nanosensor solution was quickly added, the kinetic absorbance of the mixture solution was immediately recorded by SpectraMax M3 plate reader (Molecular Devices) at the wavelength of 412 nm. The time that the solution reached the peak plateau was used for calculation of enzymatic activity via Equation (4).

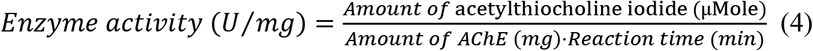

### Characterization of ACh nanosensors

The UV-Vis spectrum of DNA scaffold/sensors was measured by a NanoDrop™ 2000c Spectrophotometer (Thermo Scientific). Native PAGE was conducted to characterize DNA nanostructures by using 4% TBE gel and 1x TBE buffer at a constant voltage of 200 V (electrophoresis system Thermo Scientific) in an ice water bath. The *in vitro* reversibility test of ACh nanosensors was conducted in a slotted imaging bath (RC-21BRFS, Warner Instrument). The sensors were carefully introduced into a microdialysis hollow fiber (13 kDa, Spectrum Laboratories) via capillary action. The two ends of the dialysis tube were then sealed by super glue, and quickly attached to the glass bottom of the flow chamber. After the cap of the chamber was closed, 10 mL Ringer buffer was slowly injected into the chamber to prevent any air bubbles. The tubing was then connected to a peristaltic pump (Fisherbrand) and waste bottle. The chamber was carefully placed on the confocal imaging stage (LSM 800, Zeiss) under a 20x objective. An ACh solution of 100 μM (acetylcholine chloride, Millipore-Sigma) was injected into the chamber while the imaging started. After 2 min of imaging, the remaining ACh was washed away by Ringer’s buffer to allow the sensor to recover to the initial state. The injection of ACh was repeated 5 times.

AFM measurement was conducted following literature precedents^56^. All buffers were freshly prepared and filtered through 0.22 μm filters on the experiment day. In short, freshly cleaved mica (Ted Pella) was incubated with 20 μL of 100 m M NiCl_2_ for 1 min and rinsed with 50 mL ultrapure water. The mica was then quickly dried by filter paper, immediately followed by deposition of 20 μL of 1 nM ACh nanosensors in deposition buffer [10 mM HEPES (pH 7.5), 10 mM MgCl_2_ + 25 mM KCl]. After 2 s, the mica surface was gentle rinsed by ~10 mL of deposition buffer and 2 mL of imaging buffer [10 mM HEPES (pH 7.5), 10 mM NiCl_2_ + 25 mM KCl], in sequence. The same imaging buffer was also deposited onto the microscope tip. Samples were imaged in “ScanAsyst mode in fluid” using a Dimension FastScan microscope with PEAKFORCE-HiRs-F-A tips (Bruker Corporation)^57^.

### Animal preparation and imaging

Adult male and female C57BL/6J (RRID: IMSR_JAX: 000664) and ChAT^BAC^-eGFP (RRID: IMSR_JAX: 007902) mice between the ages of 4 to 8 weeks old were obtained from the Jackson Laboratory. All mice were maintained under the conditions mandated by Northeastern University’s Division of Laboratory Animal Medicine (DLAM). All animal procedures were approved by the Institutional Animal Care and Use Committee (IACUC) at Northeastern University and were in accordance with the National Institutes of Health guidelines. To better visualize the staining of the ACh nanosensors in the SMG, ChAT^BAC^-eGFP mice that specifically express green fluorescent protein (GFP) in the cholinergic cells and pre-synaptic axons fibers were used. In all animal experiments, the mice were anesthetized by inhalation of 1.5-2% isoflurane and placed on a rodent heating pad (Braintree Scientific, Inc.) set at 37.7°C. The mice were placed on their dorsal side under the dissection scope (Fisher Scientific) and after using depilatory cream, a small incision (about 1 cm) on the neck region was made, exposing the salivary glands. Fine tweezers (Fisher Scientific) were used to separate the salivary ducts from surrounding connective tissue. Anesthetized mice were then carefully transferred to the stage of an upright fluorescence microscope (BX61WI, Olympus) for *in vivo* imaging experiments. W-VIEW GEMINI beam splitter (Hamamatsu) was incorporated into the optical path to image the pHAb and BTX-AF647 signals simultaneously (Supplementary Fig. 11).

### In vivo microinjection of nanosensors

The nanosensors or other DNA nanostructures were administered to the SMG of living mice via microinjection. After the initial dissection work as described previously, the salivary ducts were lifted up and supported by an insulated metal platform controlled by a micromanipulator for stabilization. A borosilicate glass tube with 1.0 mm/0.78 mm OD/ID was pulled using a P-97 micropipette puller (Sutter Instrument). Fig. 3a shows the tip of a pulled needle. The pulled glass needle was loaded with 4 μL of nanosensor solution and it approached the submandibular ganglion from the proximal end at an angle of 25°using the Eppendorf InjectMen 4. Due to the use of a higher magnification water-immersion lens, vacuum grease (Dow Corning) was carefully applied around the incision area to contain an aqueous solution without leakage. Afterwards, the cavity created by the incision was filled with Ringer’s solution and then using a 63x objective, the needle could precisely pierce into the SMG and enable nanosensor injection. The nansensors were incubated within the SMG for 5 min, followed by washing with Ringer’s buffer for 10 min to remove unbound nanosensors.

### Ex vivo ACh calibration

Anesthetized mice were sacrificed by cervical dislocation immediately after the microinjection of nanosensors/control nanosensors. The SMG along with the salivary duct and the gland were quickly excised under a dissection scope. The dissected tissue was positioned within an open diamond bath imaging chamber (RC-26, Warner Instruments). The salivary duct was carefully stretched with an insect pin as a metal harp (Warner Instruments) was placed on top to suppress and stabilize the SMG (Fig. 2). After placing the chamber onto the microscope stage, 200 μL of freshly prepared Ringer’s buffer (pH=7.3-7.4) was added under the 63x water-immersion objective, followed by 500 μL injections of ACh solutions ranging in concentration from 0.001 μM to 10 mM. The final concentration of ACh was adjusted based on a dilution factor of 1.4. The entire experiment was performed using an Olympus BX61WI fluorescence microscope. After each measurement, the ACh solution was carefully removed using Kimwipes, followed by three rounds of washing with Ringer’s buffer. The sample was then immersed and incubated in the buffer for 10 min to enable the nanosensor to recover to baseline fluorescence signals. For the calibration of control nanosensors, we repeated the same procedure, with the exception of using citrate-phosphate buffers for pH<6 and phosphate buffers for pH≥7. PBS (pH=7.3-7.4) was used for intermittent washings in between measurements of the control nanosensors.

### Electrical stimulation

After microinjection of the nanosensors, a parallel bipolar electrode (FHC Inc.) was connected to a micromanipulator arm. Using an attached joystick module (Burleigh), the electrode was placed directly on the salivary duct, sitting above the SMG (closer to the lingual nerve). Electric pulses were generated by MyoPacer field stimulator (IonOptix LLC) triggered by a pulse generator (Prizmatix). Bipolar pulses with an amplitude between 3-10V, frequency of 10Hz, pulse duration of 10ms, and pulse number between 10-100 were delivered to the lingual nerves that innervate the SMG.

### Drug inhibition of ACh release

After microinjection of the nanosensors, Vesamicol (Millipore-Sigma, 100 μM in Ringer’s buffer) was added dropwise to the neck cavity and incubated for 10 min to block the release of endogenous ACh. After each measurement was imaged and recorded, the drug solution was washed away with Ringer’s buffer, and the neck region was bathed in Ringer’s buffer for another 10 min to allow for the nanosensors to return to baseline fluorescence levels before recording the next measurement.

### AQuA analysis

The AQuA software^28^ was used to accurately model and quantify the changes in intensity of nanosensor fluorescence in the SMG of a living mouse. Initially, a TIFF file of the experiment was uploaded with a set of preset values that described the frame rate, spatial resolution, and regions on the boundary of the video that were removed to apply motion correction. We defined the area to be analyzed by drawing a boundary around the entire SMG. The program determined and eliminated background noise from the video by increasing the intensity threshold scaling factor and the signal was smoothed with a Gaussian filter. Events were determined by increasing the temporal cut threshold, removing all events below the intensity threshold. These events were then grouped together into “super events” based on the distance and time difference between the single events. These events were then grouped together into “super events” based on the distance and time difference between the single events.

#### Acknowledgements

We thank funding from NIH SPARC OT2OD024909. We thank funding from NIH R01NS099254 and NSF 1604544. We thank Dr. Jeff Lichtman for early discussions on the imaging of the submandibular ganglion. We thank imaging support from the Northeastern University Institute for Chemical Imaging of Living Systems (CILS). We thank Dr. Jennifer Morales for early experimental discussion. We thank Mr. Alex Lovely for assistance on microscopy imaging. We thank Dr. Qiong Wei for assistance on AFM. We thank Mr. Qizhong Wang for assistance on data analysis.

#### Author contributions

J.R.M. and H.A.C. supervised the project. J.X., H.Y., J.R.M. and H.A.C. designed the experiment and analyzed the data. J.X. and H.Y. conducted the experiments and wrote the manuscript with input from other authors. M.M. assisted *in vivo* experiments and live imaging. K.E.P developed the AQuA software, and advised on its implementation in this study. N.M. performed propagation stimulation using AQuA.

