## Supplementary information text for "A DNA-based optical nanosensor for *in vivo* imaging of acetylcholine in the peripheral nervous system"

### Supplementary Figures

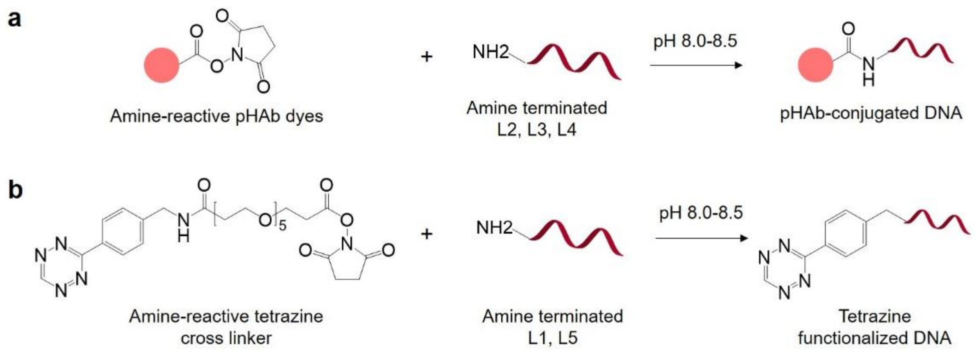


**Supplementary Fig. 1. Modification of oligonucleotides with pH-sensitive fluorophores and tetrazine functional group.**

(a) pH-sensitive fluorophores (*i.e.*, pHAb) were conjugated to strands L2, L3, and L4 via NHS ester reaction.
(b) Tetrazine functional group were linked to the strands L1 and L5 by an amine-reactive tetrazine cross linker via NHS ester reaction.

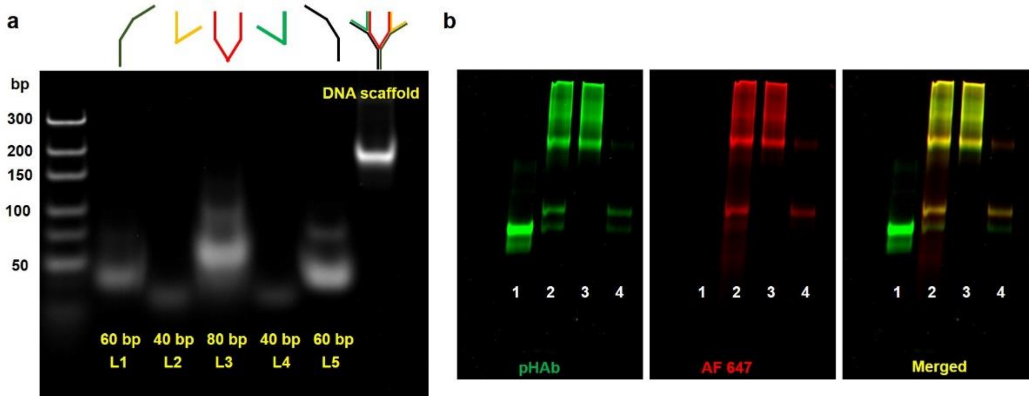


**Supplementary Fig. 2. Native PAGE electrophoresis of ACh nanosensors.**

(a) Single strands (i.e. L1 to L5) and the assembled DNA scaffold. The PAGE gel was poststained with ethidium bromide.
(b) Same PAGE gel as Figure 2e imaged under fluorescent channels of pHAb and Alexa 647. Lane 1: unpurified ACh nanosensor; Lane 2: unpurified ACh nanosensor; Lane 3: Purified ACh nanosensor (Main peak); Lane 4: impurities including AChE, DNA scaffold, BTX-conjugated DNA scaffold (Second peak).

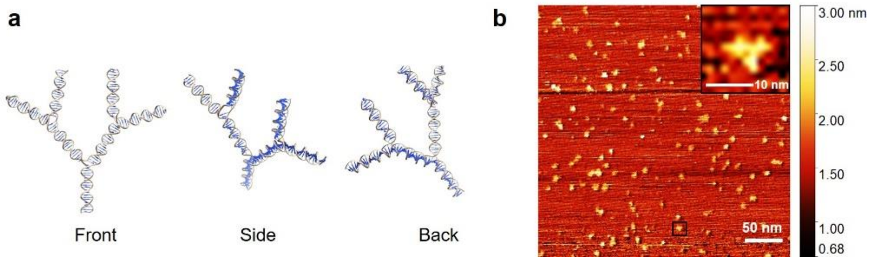


**Supplementary Fig. 3. Stimulation and AFM imaging of ACh nanosensors.**

(a) Structure of DNA scaffold stimulated by the Nanoengineer-1 Program and viewed by UCSF Chimera 1.14.
(b) AFM imaging of ACh nanosensors in liquid mode. Scale bar: 50 nm. The inset is the enlargement of the black boxed area. Scale bar: 10 nm.

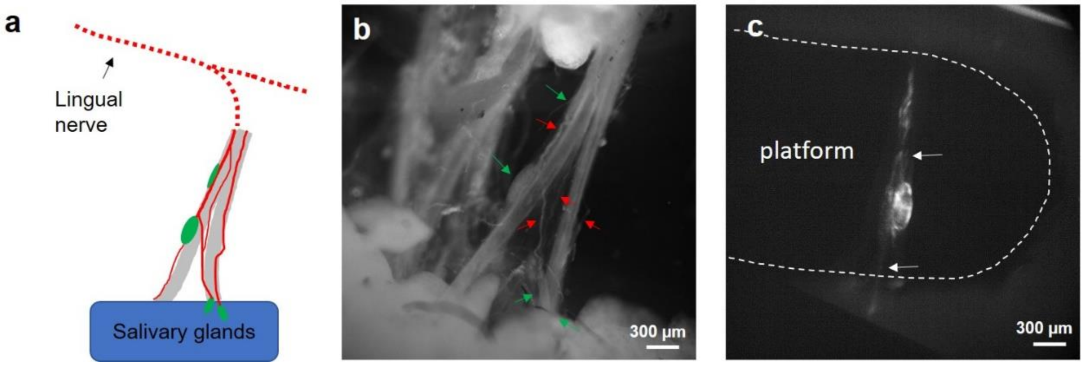


**Supplementary Fig. 4. ACh nanosensor-injected submandibular ganglion (SMG).**

(a) Schematic picture of SMGs (green) and innervating nerves (red).
(b) Corresponding microscopic image of SMGs (green arrow) and innervating nerves (red arrow).
(c) Low magnification of nanosensor-injected SMG. Arrows indicate nerve tracts entering and exiting the SMG.

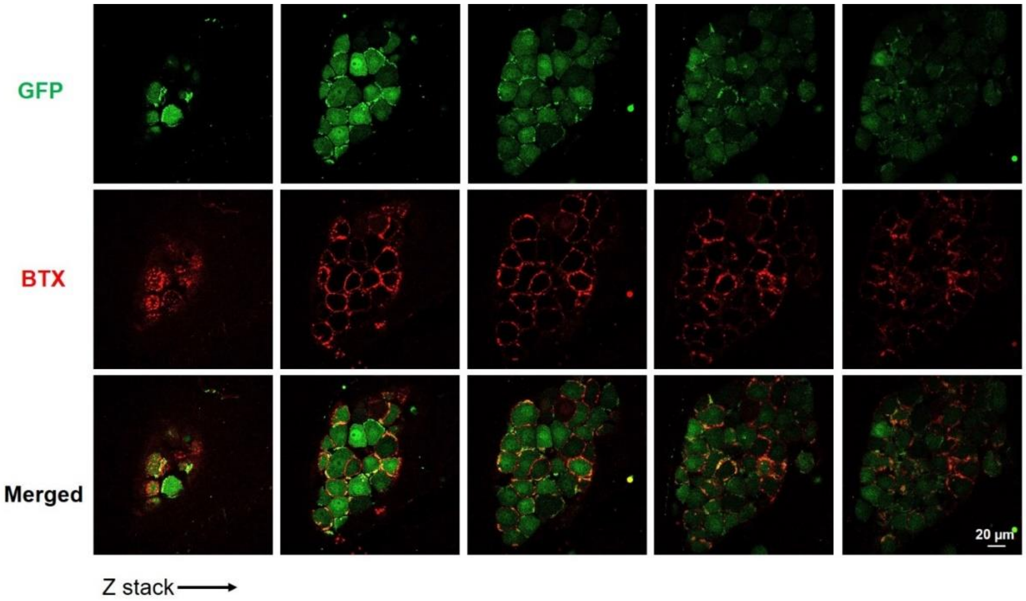


**Supplementary Fig. 5. Confocal z-stacks of submandibular ganglion (SMG) labelled by Alexa647-congujated α-bungarotoxin (BTX).**

The SMG was harvested from a ChAT(BAC)-eGFP transgenic mouse which expresses green fluorescent protein in both the pre-ganglionic nerve fibers and post-synaptic cell bodies. From top to bottom are images from the GFP channel, Alexa 647 channel (indicates BTX), and merged channels.

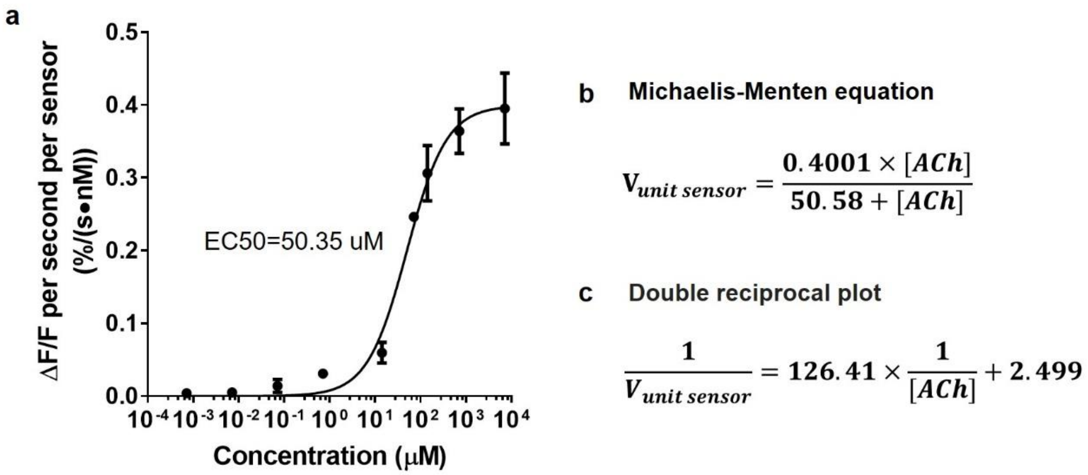


**Supplementary Fig. 6.** ***Ex vivo*** **calibration of the ACh nanosensors using unit velocity.**

(a) The sigmoidal nonlinear curve was fitted by the Michaelis-Menten equation with the constants =0.3991 %/(s·nM) and =50.35 µM (EC50). The data was replotted by dividing the sensor's initial velocity shown in Figure 4c by the nanosensor concentration. The concentration of the nanosensors was calculated based on the fluorescence intensity by a calibration curve using nanosensors of known concentrations on the same imaging microscope.

(b) Displays the fitted Michaelis-Menten equation in (a).

(c) A double reciprocal plot was derived from (b) to facilitate the quantification of endogenous ACh release.

Error bars denote S.D. resulting from triplicate experiments.

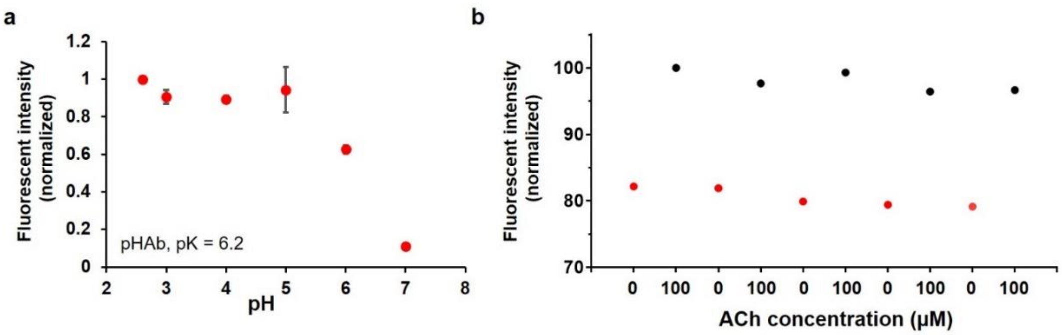


**Supplementary Fig. 7.** ***In vitro* characterization of ACh nanosensors.**

(a) Normalized fluorescence change of free pHAb fluorophores under different pH ranging from 2.5 to 7. Error bars denote S.D. resulting from triplicate experiments.

(b) Normalized fluorescence change (pHAb channel) of ACh nanosensors under five repeated injections of ACh solution ([ACh]=100 μM) in a confocal chamber.

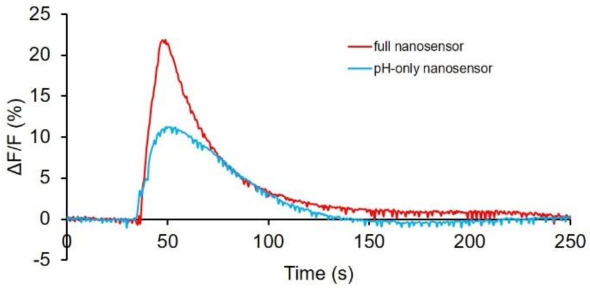


**Supplementary Fig. 8.** ***In vivo* stimulation of ACh nanosensor (full sensor, red curve) and control nanosensor (blue curve).**

Stimulation parameters are identical for both sensors: 10V, 10Hz, 10ms duration, 100 pulses.

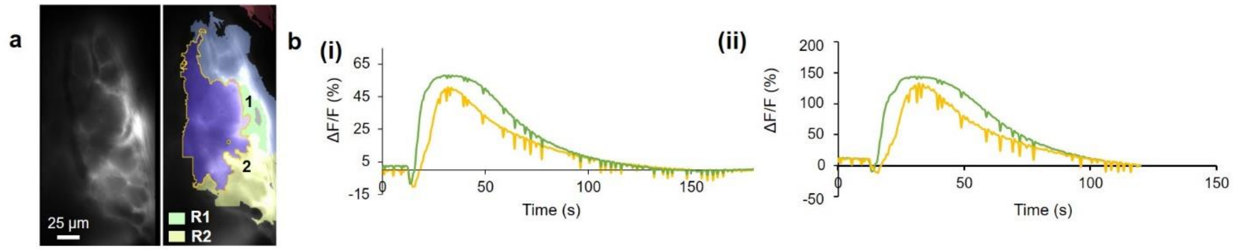


**Supplementary Fig. 9. ΔF/F of fluorescence intensity from pHAb channel in response to electrical stimulation**

(a) Two representative events were labeled as R1 and R2. The events represented groups of ACh nanosensors fluorescing at the same time and within the same region, and were automatically identified and grouped by AQuA software based on changes in intensity of individual pixels. Stimulation parameters: 8V, 10 Hz, 10 ms duration, 100 pulses.

(b) (i) Plot of ΔF/F with respect to time in the R1 and R2 in (a) based on traditional ROI-based analysis
(ii) Plot of ΔF/F with respect to time in the R1 and R2 in (a) based on AQuA analysis

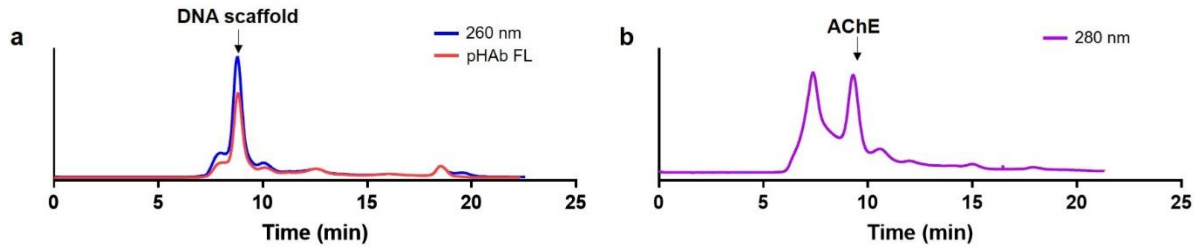


**Supplementary Fig. 10. Size exclusion HPLC purification.**

(a) DNA scaffold after annealing from 95 ºC to 4 ºC.

(b) AChE of high purity from the commercial AChE product (from eel extract).

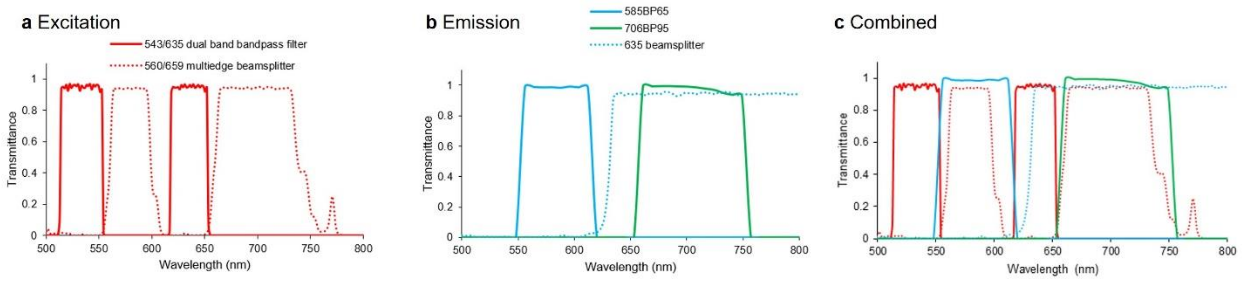


**Supplementary Fig. 11. Spectral profiles of filters and beam splitters used in the dual-color imaging mode.**

(a) Spectra of dual band bandpass filter (543/635 nm) and multi-edge beamsplitter (560/659 nm) on the excitation end of the optical path.
(b) Spectra of two bandpass filters (585/65 and 706/95 nm) and longpass dichroic (635 nm) mirror on the emission end.
(c) Combined spectra of five optical components.

Solid line: bandpass filter; dashed line: beamsplitter. Data: courtesy of Semrock, Inc.

### Supplementary Table

#### Supplementary Table 1. Oligonucleotides that assemble the DNA scaffold

Sequences (5' to 3') and the functional groups on the terminal ends of 5 oligonucleotide strands (*i.e.*, L1, L2, L3, L4, L5) used in the DNA scaffold.

| Strands | Sequence (5' to 3') and functional groups |
|---|---|
| **L1** | [Amine]TCTGAAAGTACTGACGAGCTAACATGGCTGCGGCAGAATCCCTCACTATGCGAGTTGACC[Thiol] |
| **L2** | [Amine]GGTCAACTCGCATAGTGAGGGAGTCGTGAGTACTAATAGT[Thiol] |
| **L3** | [Amine]ACTATTAGTACTCACGACTCGATTCTGCCGCAGCCATGTTTCGCCAGAATGCCAGTCAGCATTAAGGAGAGCTCAGGGCA[Amine] |
| **L4** | [Thiol]TGCCCTGAGCTCTCCTTAATAGCCTACATCCTACCAGAGG[Amine] |
| **L5** | [Thiol]CCTCTGGTAGGATGTAGGCTGCTGACTGGCATTCTGGCGAAGCTCGTCAGTACTTTCAGA[Amine] |

### Supplementary Video

**Supplementary Video 1.** Microinjection of ACh nanosensors into SMG via a glass needle.

**Supplementary Video 2.** Ex vivo calibration in ACh nanosensor-injected SMG.

**Supplementary Video 3.** The response of ACh nanosensors in SMG upon 8V stimulation and the corresponding AQuA analysis.

**Supplementary Video 4.** AQuA analysis of the response of ACh nanosensors in SMG upon 3V stimulation.
