## Supplementary figures and images for "A DNA-based optical nanosensor for *in vivo* imaging of acetylcholine in the peripheral nervous system"

### Supplementary information Video 1

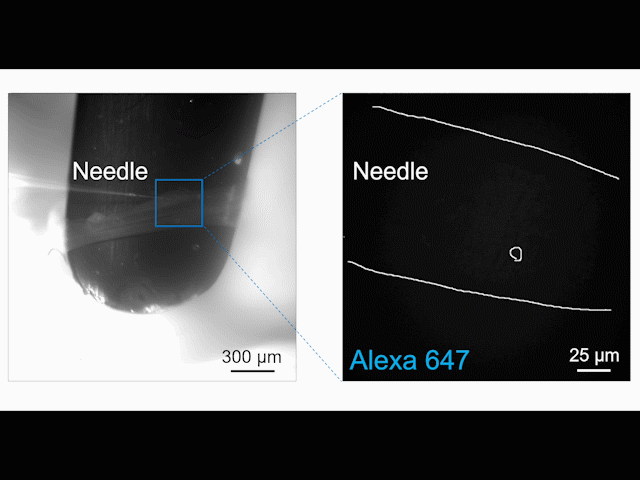

### Supplementary information Video 2

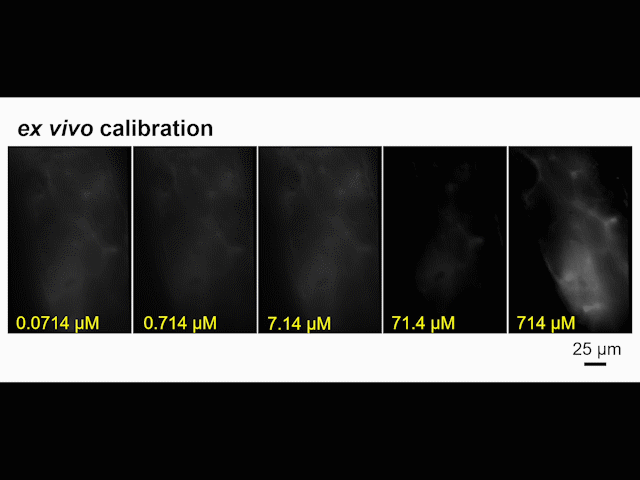

### Supplementary information Video 3

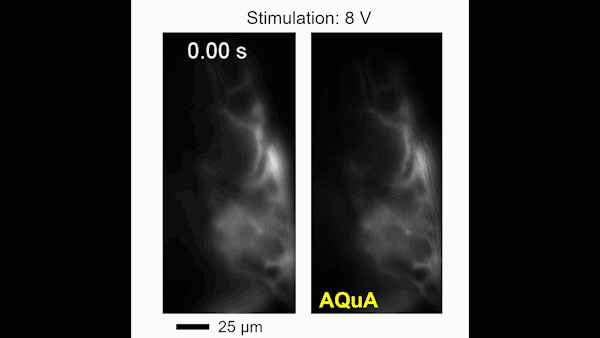

### Supplementary information Video 4

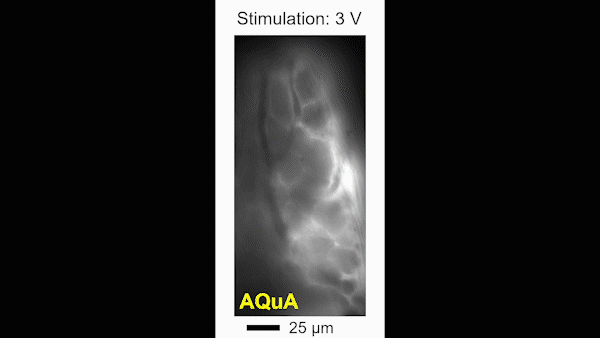
